# Fully Convolutional Networks for pathophysiological Intracranial Pressure waveform classification

**DOI:** 10.1101/2020.11.17.381517

**Authors:** O. Vrabie, R. Faltermeier, N.O. Schmidt, A. Brawanski, E. W. Lang

## Abstract

It is generally assumed that the analysis of intracranial pressure (ICP) waveforms could be used for the detection of multiple cerebral pathophysiologies. A main obstacle for the analysis of ICP waveforms is given by the large variation of their generating signal, the arterial blood pressure (ABP). Using extended principal component analysis (PCA) we show that it is possible to distinguish between ICP waveforms generated by pathological ABP waveforms, e.g. in the case of a heart failure, without loosing information about the state of the cerebral compliance. We also create a dataset for ICP pulse classification that can be used to train models to distinguish between an intact and diminished cerebral compliance, as a function of the ICP-generating ABP pulse. As a baseline for classification we build a fully convolutional network (FCN) that reaches high performance with relatively few data.

## 1 Introduction

The treatment of patients suffering from Traumatic Brain Injury (TBI) or subarachnoid hemorrhage (SAB) aims to avoid secondary brain injury by continuous monitoring of different physiological parameters like intracranial pressure (ICP) and arterial blood pressure (ABP). During treatment a main focus is placed on the assessment of the cerebral compliance which can be defined as the ability of the cerebral compartment to compensate for volume changes in terms of pressure. A diminished cerebral compliance caused by a severe brain swelling may lead to pronounced increase in ICP and subsequently to a life threatening reduction of the cerebral perfusion.

From a time series analysis point of view a diminished cerebral compliance should be detectable by means of specific patterns occurring in the ICP recording during an episode of severe brain swelling. A common method to identify such patterns is the examination of individual ICP waveforms. ICP waveforms are intracranial pressure waves that are generated by the ABP pulse waves running through the cerebral vessels and are strongly influenced by the state of the cerebral compartment. However evaluating ICP waveforms in real time by a human expert is a demanding task simply due to the mere amount of data.

In this regard considerable efforts have been spent to automatize both extraction and categorization tasks. One prominent example is the morphological clustering and analysis of intracranial pressure (MOCAIP) algorithm [9, 10]. The algorithm first identifies an individual ICP pulse wave and then computes a set of 24 so-called MOCAIP metrics to characterize this individual pulse. Among other things the metrics contain the height of the different peaks of such a pulse and the distance between them. Subsequently the metrics are fed into a separate classifier. However, the 24 manually picked metrics contain redundant information and quickly become very time-consuming to be analyzed.

Contrary, some efforts are focused on leaving the entire feature extraction procedure to a non-biased “agent”. An example of this paradigm is presented in [18] where cerebrospinal fluid pulse pressure waveforms (CSFPPWs) are classified by an artificial neural network (ANN). The authors defined four distinct morphological classes of the cerebrospinal fluid (CSF) pulses ranging from normal to pathological and subsequently labeled 486 CSFPPWs from 160 patients with different neurosurgical conditions, to form the training set of the ANN. The normalized waves were mapped onto an embedding vector of ten radial basis function coefficients. This vector served as input for the ANN. After training the network, 60 new CSFPPWs were used to evaluate it’s accuracy by comparing the performance of the model with a human expert. They reached 88.3% classification concordance, an unquestionably good result, considering that the model was exposed to less than 500 examples.

The influence of human experts happens either at the feature definition and selection phases, or during picking and labeling the training examples. Safely excluding the human expert from the analysis can be an almost impossible task but would help to bypass the above mentioned issues. It could be done in phases, as was demonstrated in the automatic feature selection for the MOCAIP algorithm [9], letting the model to find patterns on its own using, for example, an ANN-based classifier like in Nucci *et al.* [18], or combining them towards a more generalizable approach.

In this study we analyze the influence of diminished cerebral compliance on the ICP waveform using a fully convolutional network (FCN). To minimize the influence of human expert knowledge on the classification task, we made use of a mathematical toolkit called Selected Correlation Analysis (SCA) that reliably detects episodes of diminished cerebral compliance by analyzing the coherence of low frequency components of ABP and ICP signals [7, 8, 19]. To identify morphological categories of waveforms for the above mentioned episodes we use principal component analysis (PCA) and several clustering methods. Additionally we determined the influence of the ABP waveform on the ICP wave form using a similar procedure. First we looked for three different types of ABP waveforms, described in the literature [6, 17], representing a healthy heart, a heart dysfunction and a severe sepsis that already influences the stiffness of the vessels. With a further cluster analysis we find that the above mentioned morphological categories of ICP waveforms can be additionally subdivided when accounting for the morphology of the associated ABP waveforms. ICP pulses could be separated by means of both the state of compliance and the type of the input ABP waveform. The FCN model was trained on the six-class dataset (two compliance states *×* three ABP waveform states) reaching a high accuracy on a hold-out test set. In addition to high classification accuracy, the use of an FCN allowed us to construct class activation maps (CAMs), which helped visualizing exclusive regions in the ICP pulses that determined the pulse classification.

## 2 Materials and Methods

### 2.1 Patients and data acquisition

The data was collected from a cohort consisting of seven patients suffering from TBI and Subarachnoid hemorrhage (SAH). The summary of patients’ physiological variables is depicted in Table 1 (for more details see Table 1). The ICP time series were recorded continuously in the intensive care unite (ICU) using an intraparenchymal piezoelectric sensor, which was inserted into the brain tissue at a depth of around 2 *cm*. The ABP time series were simultaneously recorded at the radial artery. Both quantities were measured continuously with a sampling rate of 1 *kHz* [8, 13, 21].

**Table 1:**
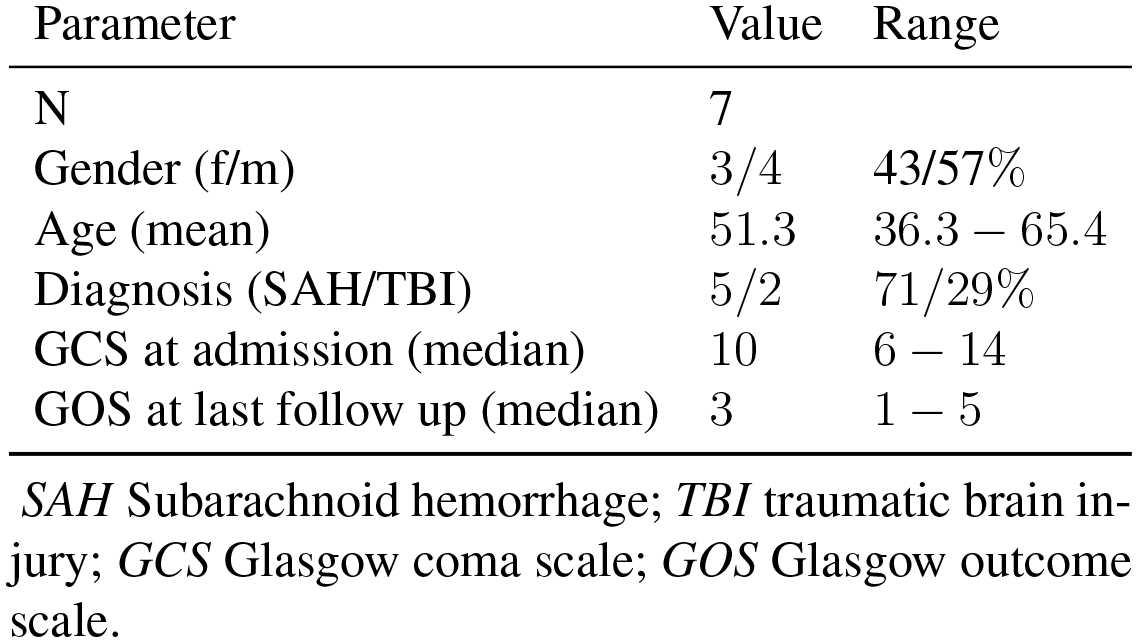
Mean and median of physiological variables Parameter Value Range

The study was conducted in accordance with the ethical guidelines of the Institutional Review Board of the University of Regensburg. Informed consent was obtained from the patient’s relatives and all study results were stored and analysed in an anonymized fashion.

### 2.2 Selected Correlation Analysis

The motivation for the development of the selected correlation analysis (SCA) comes from predictions of a mathematical model of cerebral perfusion and oxygen supply [3, 8]. Simulating the time evolution of ICP under the condition of a diminished cerebral compliance caused by a severe brain swelling, the model predicts two different states. In case of an intact cerebral autoregulation the diminished compliance leads to a negative correlation between the homeostatic components of the ABP and the ICP signal. In case of a disturbed autoregulation the diminished compliance leads to a positive correlation between the above mentioned signals. If autoregulation is intact and the compliance is not disturbed, no such correlations are predicted.

To detect the above mentioned correlations in real world data we use SCA, a set of mathematical tools based on frequency analysis in a time-resolved way [8]. By splitting both time series into isochronous, fixed length segments, the multitaper coherence spectrum and the multi-taper power spectra of the segments are calculated. From this spectral information an index called selected correlation (sc) is deduced, which reflects the strength of correlation of the data segments with respect to a specific frequency window. Additionally, the mean Hilbert phase difference of the segments is calculated to reflect the phasing of the data segments. Using a specific statistical test against false positives the data segments are subsequently labeled as sc positive (scp), sc negative (scn) or no sc (nsc). A positive selected correlation (scp) indicates a diminished compliance with defective autoregulation, whereas a negative selected correlation (scn) indicates a diminished cerebral compliance with an intact cerebral autoregulation. Finally, no significant correlation (nsc) indicates no disturbance of cerebral compliance or autoregulation.

In the current study, the SCA was used to determine data segments where a diminished cerebral compliance caused by a severe brain swelling is indicated (labeled scp). Additionally we determined data segments where no disturbance of the cerebral compliance is detectable (labeled nsc). Only scp- and nsc-marked segments were selected from the recordings in order to ease interpretability. The complete parameter set used for SCA was adopted from Faltermeier *et al.* [7]. In the following section, the utilization of a pre-processing routine is described.

### 2.3 Pre-processing of pulsatile data

Utilizing a Matlab package developed by Achhammer [1], representative pulses were extracted. This was accomplished by averaging over a 10 *s*-long window of SCA-catalogued ICP segments. In the first step, the routine is cutting each pulse from minimum to minimum (Fig. 1 panel B). In the next step, a pair-wise comparison between each of the raw pulses is performed, using the Jensen-Shannon Distance (JSD) as a similarity measure. The pulse with the highest mean similarity is considered a temporary reference pulse in the next intermediate step. A JSD-similarity between the reference and each of the other pulses is then computed and compared to a custom threshold - all the pulses with a value above that threshold are rejected, otherwise sent to the next step. This step prevents highly dissimilar pulses, e.g. mechanical noise or movement artifacts, from smearing the representative pulse.

**Figure 1:**
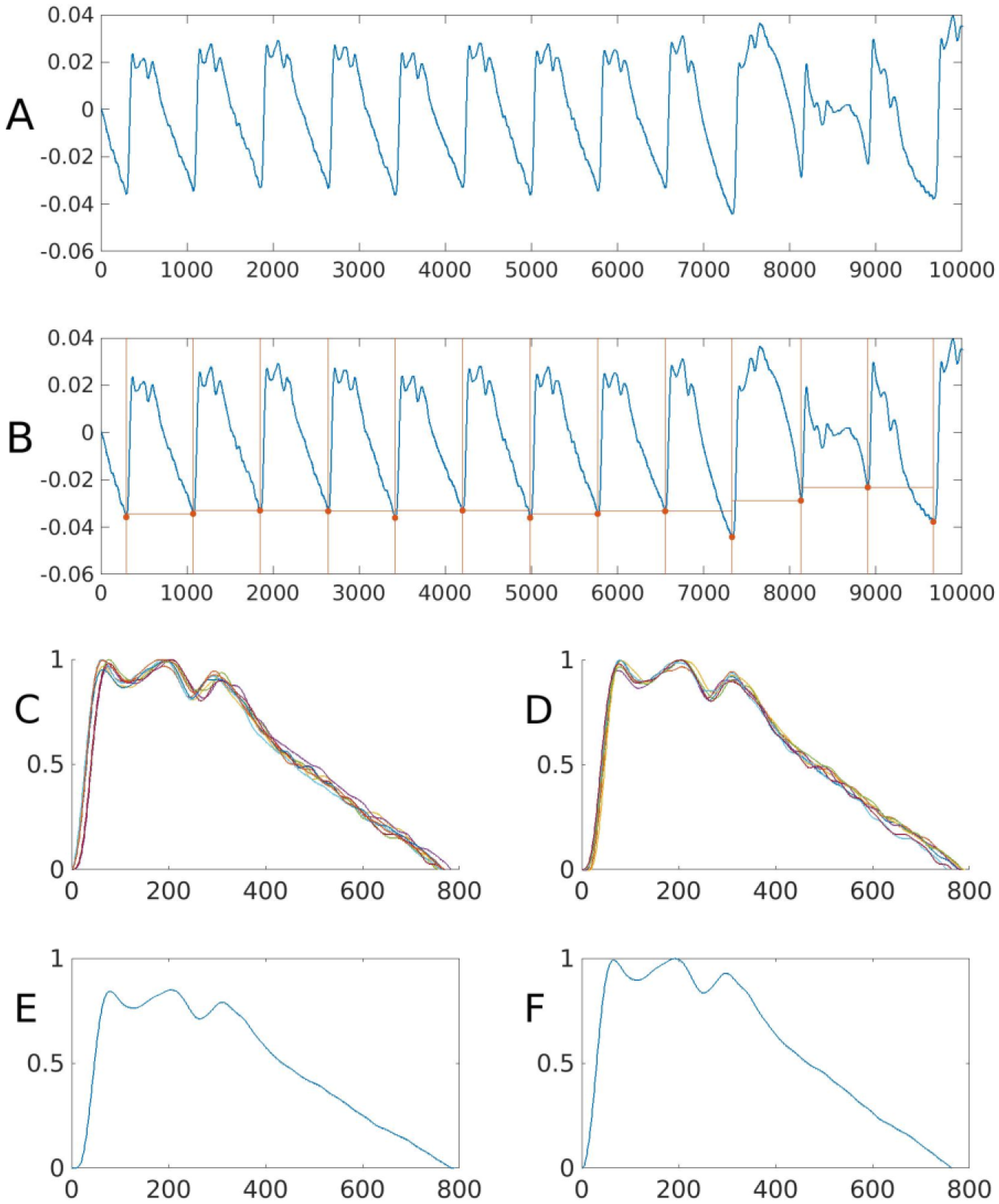
The process of creating a representative ICP pulse wave from a 10*s*- long window. (**A**) The raw sequence is cut into pieces, (**B**) from minimum to minimum, and then stacked together (**C**). In this example the last three pulses from **B** were excluded by the JSD-based artifact removal. The curves are shifted along the *x*-axis s.t. the Euclidean distance between pulses gets minimal (**D**). The resulted stack of waves is averaged (**E**) and normalized (**F**). On the *x*-axis the time is captured in milliseconds. On the *y*-axis (**A-B**) the ICP amplitude is captured in mmHg, whereas in the following panels is the normalized amplitude. Figure obtained from Achhammer [1].

This is followed by the alignment of pulses by shifting the curves in order to minimize the Euclidean distance between them (Fig. 1 panels C-D). The mean of the aligned pulses is computed (Fig. 1E), and the resulting prototype pulse is normalized (Fig. 1F). An additional step was included in order to ease the further analysis, by resizing the pulses to an average length of 780 sample points.

### 2.4 Extracting pulse modes

#### 2.4.1 PCA

In order to visualize the structure of the our data, the principal component analysis (PCA) was used as a means of dimensionality reduction. Given an input set, this technique performs a transformation to a new space of uncorrelated variables, called principal components (PCs), that contain most of the variance present in the original variables [12]. We applied PCA to the dataset of extracted ICP pulses from the entire cohort.

### 2.5 Clustering

Two clustering techniques were used to reveal the hidden structure of the dataset: Gaussian Mixture Models (GMMs) and HDBSCAN, both chosen for their flexibility and highly customizable parameters. GMMs represent a probabilistic method that attempts to find a mixture of multi-dimensional Gaussian probability distributions that best fit a dataset. The parameter search for these models is usually based on a maximum likelihood estimation (MLE), using the Expectation-Maximization (EM) algorithm [2]. The advantage of using GMMs for clustering comes from the ability of the user to define the number of components into which to decompose the dataset. HDBSCAN, on the other hand, is an advanced density-based clustering algorithm based on hierarchical density estimates [4, 16]. The main advantage of using HDBSCAN is that it allows clusters with varying densities of points.

#### 2.5.1 Fully Convolutional Network for time-series classification

For our purposes, we used a variation of CNNs called Fully Convolutional Neural Networks (FCN) that has been recently popularized for semantic segmentation [15]. More importantly for us, it recently achieved state-of-the-art performance from scratch in time series classification [20]. Apart from being easy to train, FCNs offer additional significant advantages like, for example, the invariance in the number of parameters in convolutional layers. The difference between a typical CNN and an FCN rests on two main points: (1) there are no local pooling or down-sampling layers in between convolutions and (2) the number of parameters is considerably lower due to the replacement of the final fully connected (FC) layer with a global average pooling (GAP) operation [14].

GAP layers were first proposed by Lin *et al.* [14] as an alternative to traditional FC layers. Instead of creating a further downsampled feature map, it takes the feature maps of size (*height × width × n*_*channels*) of the last convolutional layer and computes a single (average) number per channel resulting in (1 *×* 1 *× n*_*channels*). This procedure enforces a correspondences between feature maps and categories without the need of additional learnable parameters to be optimized, reducing therefore the risk of overfitting.

Besides the afore mentioned advantages of GAPs, they also allow building class activation maps (CAMs), which highlight regions in the input contributing to the classification, therefore aiding interpretability of the model [11, 20, 22]. In principle, they work as follows: specific units in a convolution layer are selectively activated depending on the visual patterns that fall in its receptive field. These activations are further up-sampled to match the size of the input image, constructing the CAM. See the complete definition of CAMs in 7.3.

The architecture of our model is depicted in Fig. 2. It was implemented in Keras [5] and was inspired by the FCN model from Ismail Fawaz *et al.* [11]. The one-dimensional CAMs that have been used in this study were introduced by Wang *et al.* [20]. This allowed us to visualize which regions in the ICP pulses contributed the most to the label assigned by our FCN during testing (Figs. S6 and S7).

**Figure 2:**
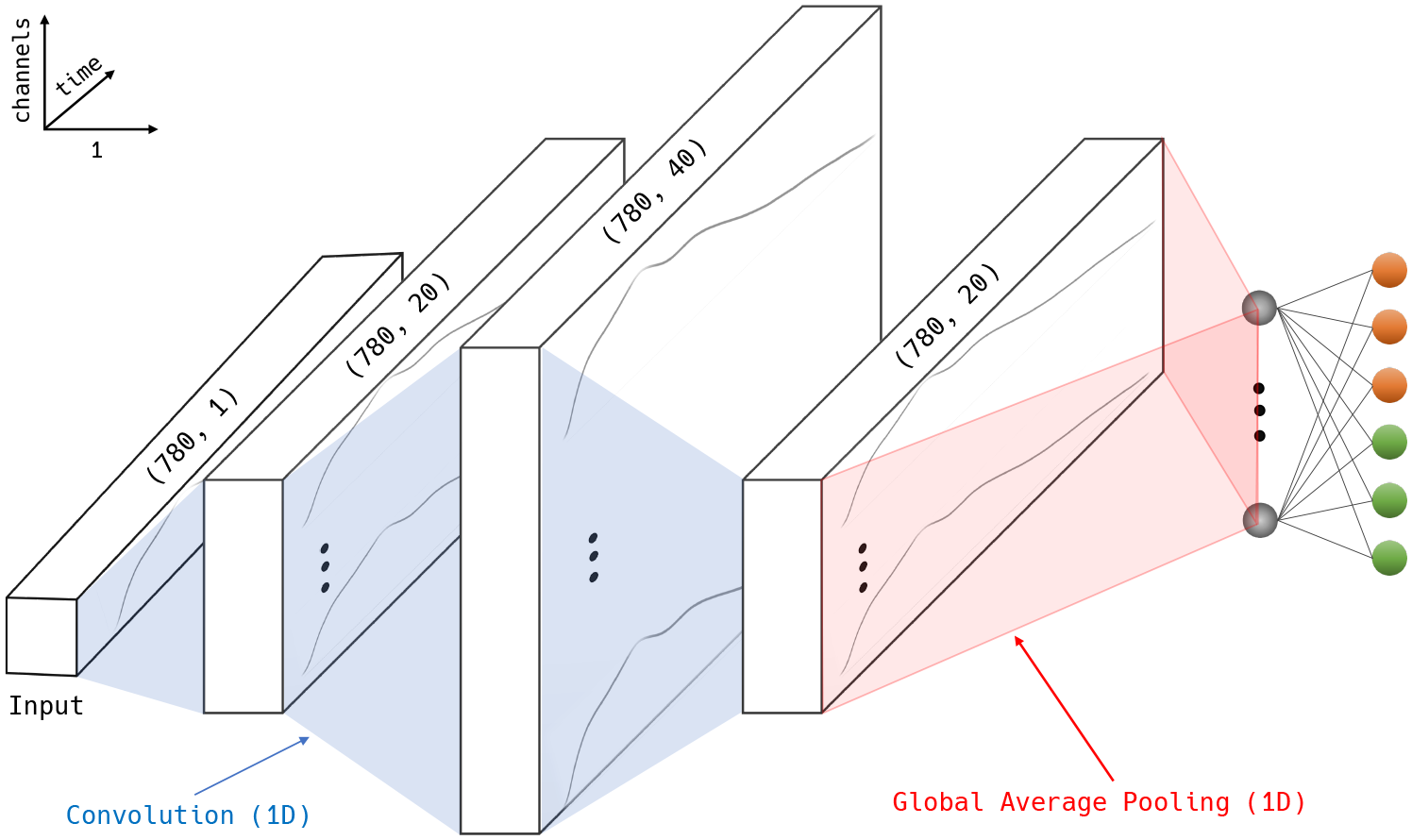
Schematic of the FCN architecture. The 1D input is subjected to three successive convolutions. The weights of the last convolutional layer undergo GAP. This is followed by the last, fully-connected `softmax` layer with 6 nodes, each representing a class (orange for scp; green for nsc).

## 3 Results and Discussion

### 3.1 Inter-patient ICP pulse classification

#### 3.1.1 Individual ICP pulses

The first step in the ICP classification pipeline was to extract individual ICP pulses using the pre-processing pipeline described in Section 2.3. As a result, we ended up with a database (Table 2) consisting of normalized scp and nsc ICP pulses from the recordings of seven human subjects.

**Table 2:**
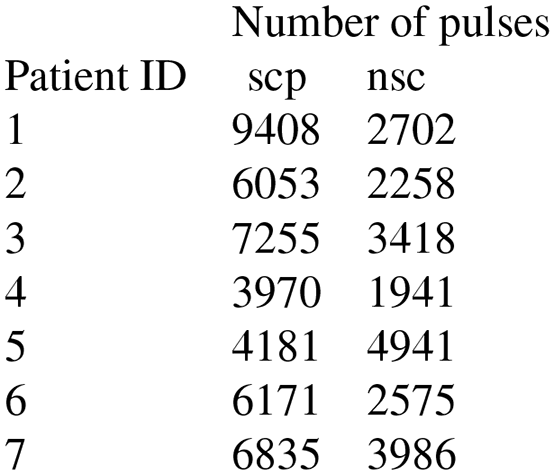
Number of pulses extracted from raw ABP and ICP time-series for each human subject.

| Patient ID | Number of pulses |  |
| --- | --- | --- |
|  | scp | nsc |
| 1 | 9408 | 2702 |
| 2 | 6053 | 2258 |
| 3 | 7255 | 3418 |
| 4 | 3970 | 1941 |
| 5 | 4181 | 4941 |
| 6 | 6171 | 2575 |
| 7 | 6835 | 3986 |

#### 3.1.2 ICP modes

After extracting all individual pulses, the second step was to identify modes, or patterns of vibration across these pulses. We achieved it by projecting pulses onto their first three principal components using PCA. The projection in Fig. 3 of the entire ICP dataset onto the first three PCs revealed that there is no major structural difference that could be responsible for the variability in pulse shape of nsc and scp data. Therefore we proceeded to a patient-wise extraction of modes by means of clustering.

**Figure 3:**
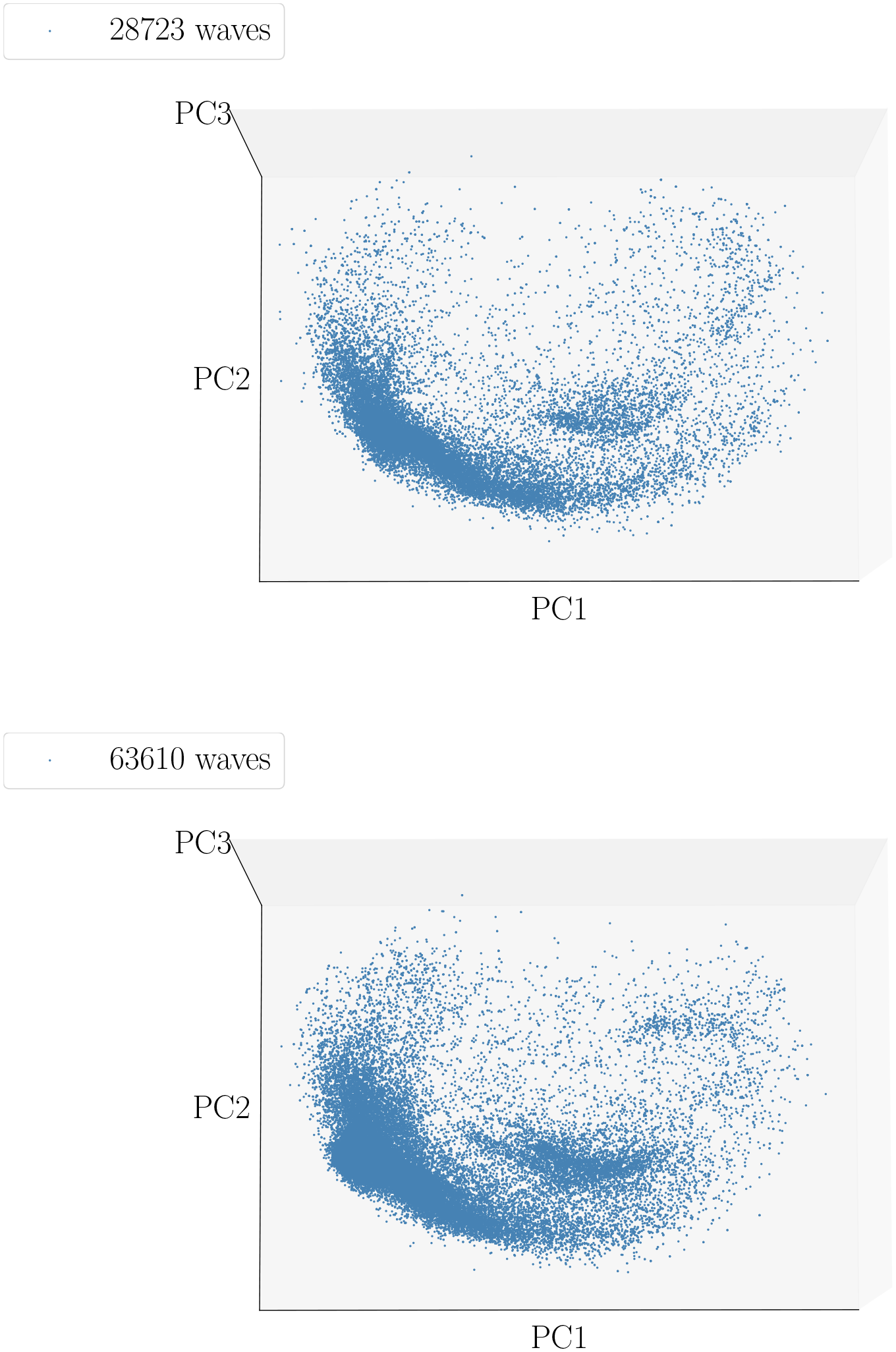
Projections of concatenated nsc (a) and scp (b) ICP pulses from all the seven patients onto first three PCs.

The results of the patient-wise clustering of pulsatile ICP are presented in grid plots in Fig. S1 showing the mean ICP pulse wave for each found cluster. Surprisingly, both scp and nsc segments amounted to almost the same number of clusters, i.e. 46 for nsc and 45 for scp. Apart from regular ICP modes there are plenty of noise modes in the data. Some of them may correspond to movements of the patient lying in bed, others could catch the plugging and unplugging of the sensors. There are also some modes that correspond to improperly cut pulses during data pre-processing.

#### 3.1.3 ICP pulse classification

The premise of this part of the study was that inspecting data from seven different patients will reveal injured brain states that could potentially capture the transitions between “ignorable” and “dangerous” conditions. Projecting ICP pulses from additional patients unveiled, as anticipated, a wide variety of waves residing in agglomerated clusters. At this point we have used the knowledge of a human expert to pick representative ICP waveforms out of the above mentioned cluster means (see Fig. S1) that best represent four classes: **nsc**, **scp**, **intermediate** and **noisy**.

Based on these classes we defined an inter-patient dataset that could be used to train a pulse classifier. As a choice for the classification algorithm we selected a Fully Convolutional Neural Network (FCN) for its performance and ease of training (see Section 2.5.1). The model was trained on a set of 42711 waves split into four classes with the following proportions: 5018 : 21946 : 4486 : 11261 (nsc, scp, intermediate and noise, respectively). After a small amount of training time, the model converged to a reasonable validation accuracy. The final test was performed on a hold-out test set, consisting of 10678 waves with class proportions: 1274 : 5483 : 1193 : 2728. This model reached 93.1 % accuracy on this set. Training and testing data sets were obtained by randomly splitting data of all seven patients.

### 3.2 Improved ICP-ABP classification

#### 3.2.1 ABP modes

In theory, the most prominent influence on the shape of the ICP wave is given by the appropriate ABP pulse wave, which can be interpreted as the generating signal of the ICP wave. Therefore physiologically-conditioned variants of the ABP waveform may have strong ramifications on the interpretation of ICP waveforms with respect to cerebral compliance. Medical literature roughly describes three systemic variants ot the ABP pulse waves, a typical healthy one, one originating from a heart failure [6] and a third one showing multiple reflections [17] due to strong vasoconstriction (see prototypes of these waveforms in Fig. 4). To estimate the influence of the above mentioned ABP pulse wave variants on the detection of scp and nsc ICP waves we first performed a patient-wise pre-processing of the ABP pulses, identical to that used for ICP pre-processing (see Section 2.3).

**Figure 4:**
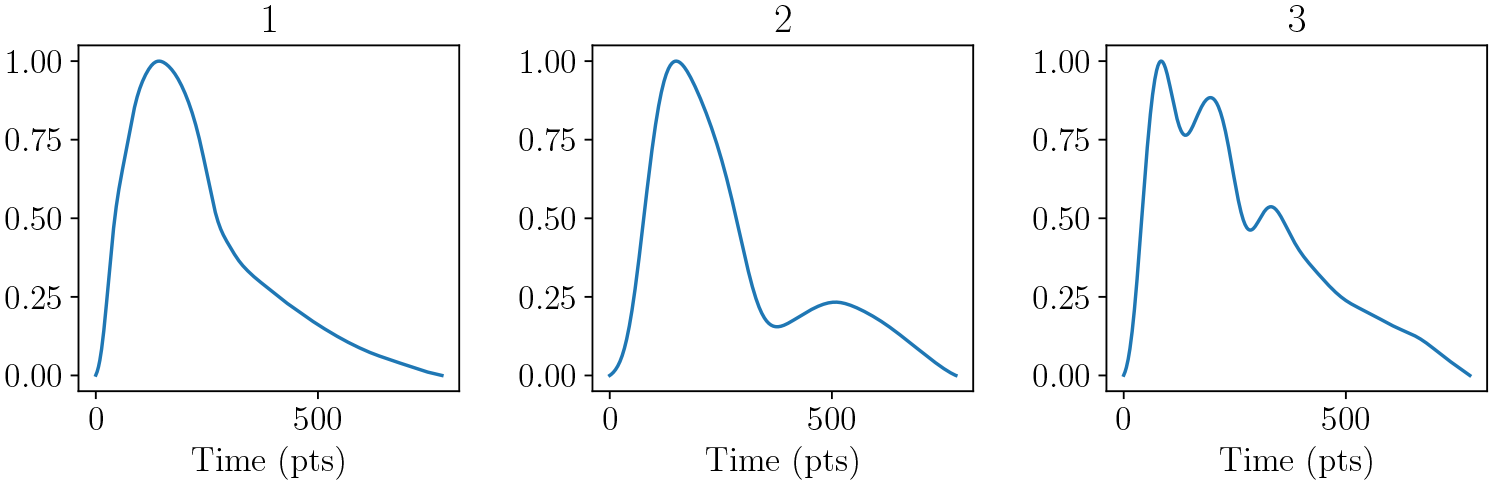
Chosen ABP WFs from our recordings. First representative WF is taken from patient 7 nsc and corresponds to a normal (most common in our cohort) ABP wave; second wave is from patient 10 scp and corresponds to a heart failure; the third comes from patient 37 scp and corresponds to multiple reflections, i.e. vasoconstriction or stiff blood vessels.

The result of the patient- and selected correlation-wise clustering of pulsatile ABP is presented in Figs. S2 and S3. At first glance, each patient exhibits at least six different morphologies for both scp and nsc types. Few noise components, e.g. nsc components 16, 20, 30 and 44 (Fig. S2) and scp components 11, 37, 44, 49 and 54 (Fig. S3), were probably caused by external mechanical perturbations. Also moderate intra-subject variation can be recognized, which, however, is stronger when comparing pulses between patients. Another aspect to be mentioned is the larger number of scp clusters in the ABP data, namely 75 in comparison with 61 nsc clusters, resulting in a significantly imbalanced dataset. On the other hand, ABP recordings contained twice as much scp than nsc annotations.

Through visual inspection, we identified all ABP clusters with cluster prototype waveforms (see Figs. S2 and S3) similar to the waveforms given in Fig. 4 as a function of sc-index. The selected ABP components are listed in Table 3.

**Table 3:** The chosen ABP components from figures S3 and S2 divided into three physiological categories: regular, heart failure and multiple reflections, and two cerebral compliance states: scp and nsc. Notation: index of the ABP component from each sc-index.

| sc-index | regular | heart failure | reflections |
| --- | --- | --- | --- |
| scp | 1, 9, 17, 45, 50 | 7, 15 | 65, 8 |
| nsc | 5, 6, 36, 46 | 17 | 31, 59 |

In order to visualize the shape of ICP pulses corresponding to chosen ABP components, we aggregated ABP pulses by physiological category and sc-index from the selected clusters and extracted the corresponding ICP pulses (Fig. 5). Furthermore, to analyze the variety of ICP pulses that contributed to those means, we turned back to PCA. The first three PCs were extracted and used to project the pulses for a further clustering step. This task was achieved using the HDBSCAN algorithm (See Fig. 6 for scp and Fig. 7 for nsc).

**Figure 5:**
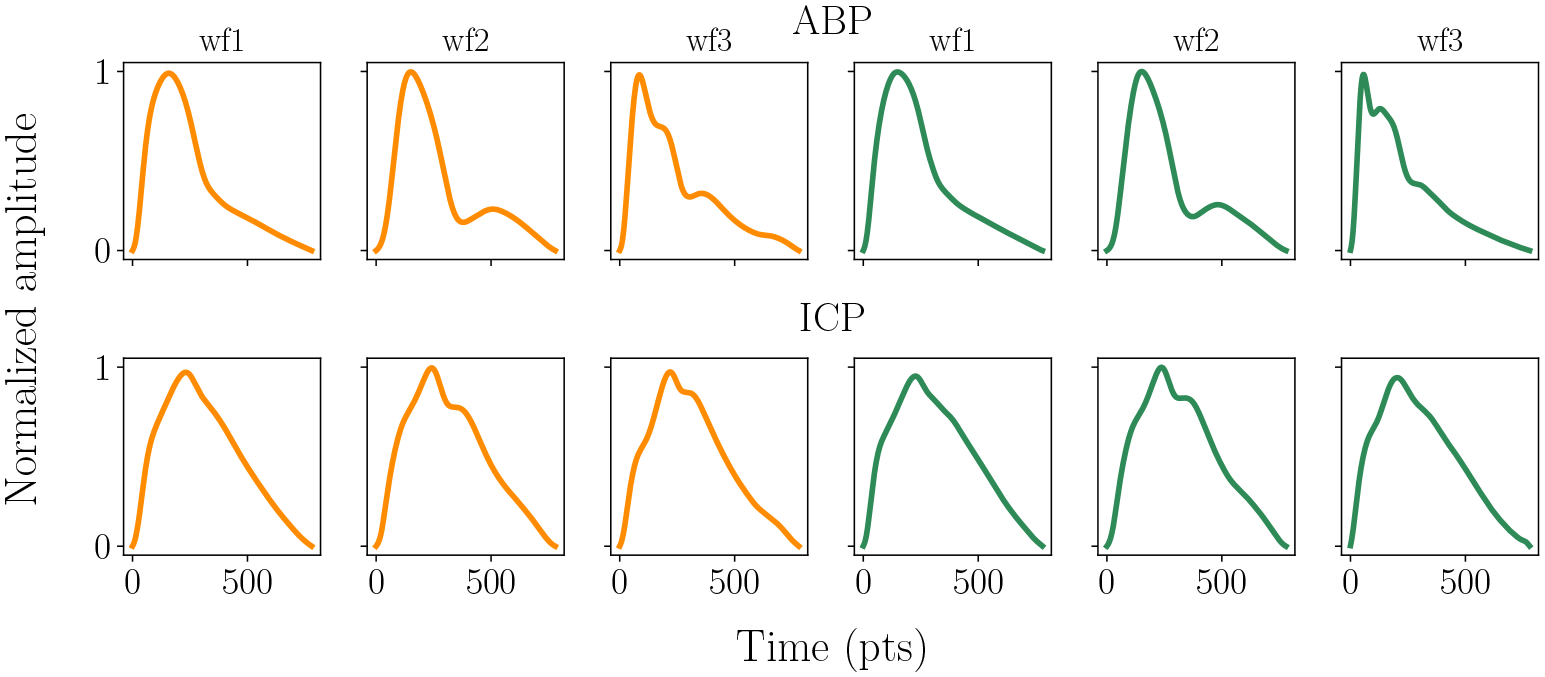
Mean of selected ABP pulses (first row) and corresponding to them ICP pulses (second row). First column corresponds to common ABP WF; second column – heart failure; third column – multiple reflections. Orange pulses corresponds to scp whereas green ones to nsc.

**Figure 6:**
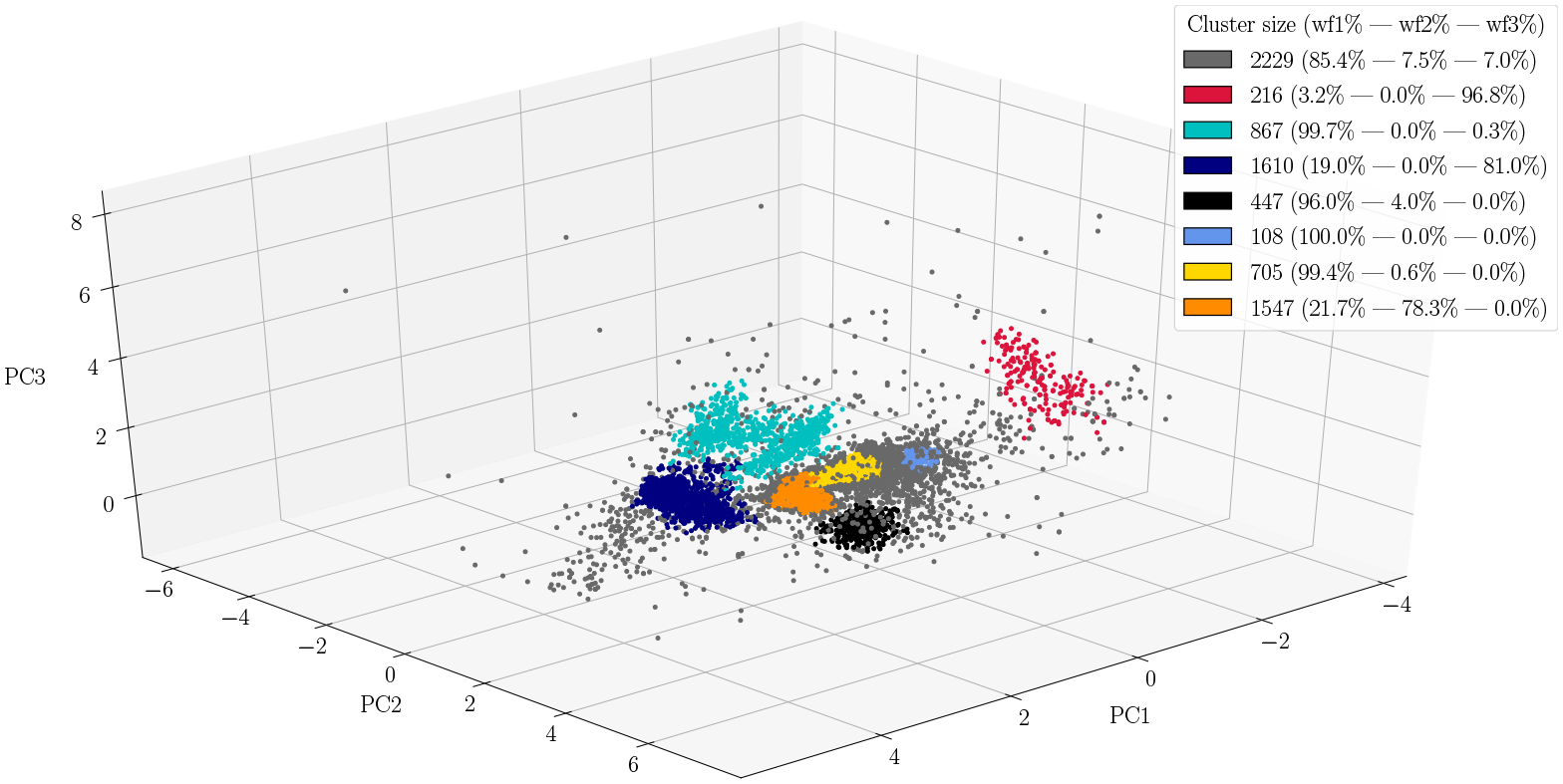
PCA-projection of **scp** ICP waves corresponding to chosen WFs. Clusters identified by HDBSCAN are marked by different colours. The gray points correspond to noise points considered by the algorithm. In the legend are captured the ratios of each WF that contributed to the found cluster.

**Figure 7:**
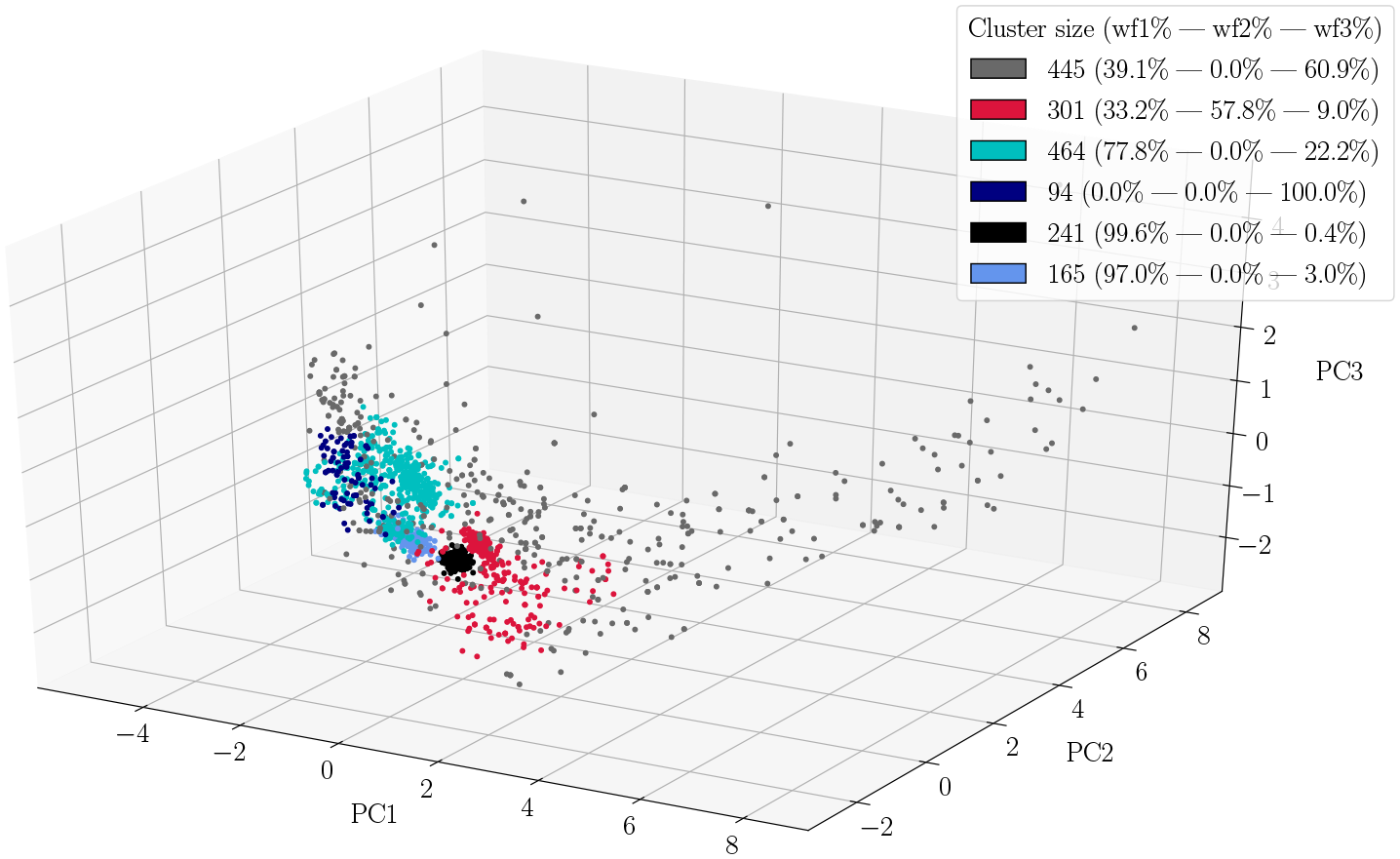
PCA-projection of **nsc** ICP waves corresponding to chosen WFs. Clusters identified by HDBSCAN are marked by different colours. The gray points correspond to noise points considered by the algorithm. In the legend are captured the ratios of each WF that contributed to the found cluster.

#### 3.2.2 Selected correlation or input ABP

By calculating the ratios of ABP waveforms (*wf1, wf2 and wf3*) that constituted each ICP cluster in Figs. 6 and 7, we could identify which of the ABP components was mainly responsible for the resulting ICP. Given these ratios, the question arose: *Are we able to distinguish between ICP waves created by the same type of input ABP waveform, but belonging to a different selected correlation type?* In other words, is the difference between selected correlation types dominating the influence of the input ABP waveform or not? To answer this question, we grouped together identified ICP clusters from both sc types to see (1) if these clusters are at all separable or, if not completely, (2) what is the rate of pulses from each sc type that build up those additional clusters.

An example of such a grouping is depicted in Fig. S4, where the 867 **scp** ICP pulse clusters and the 241 **nsc** ICP pulse clusters are both predominantly (99.7 % and 99.6 %) caused by ABP waveform 1. A GMM with 5 components was able to separate the clusters. The prototype components (cluster means) are plotted in Fig. S5.

Additionally projecting the cluster components onto the first two principal components (PCs) revealed, without exception, that all components were separable in the PC subspace using GMM clustering. All results of this pair-wise analysis are presented in Table 4.

**Table 4:**
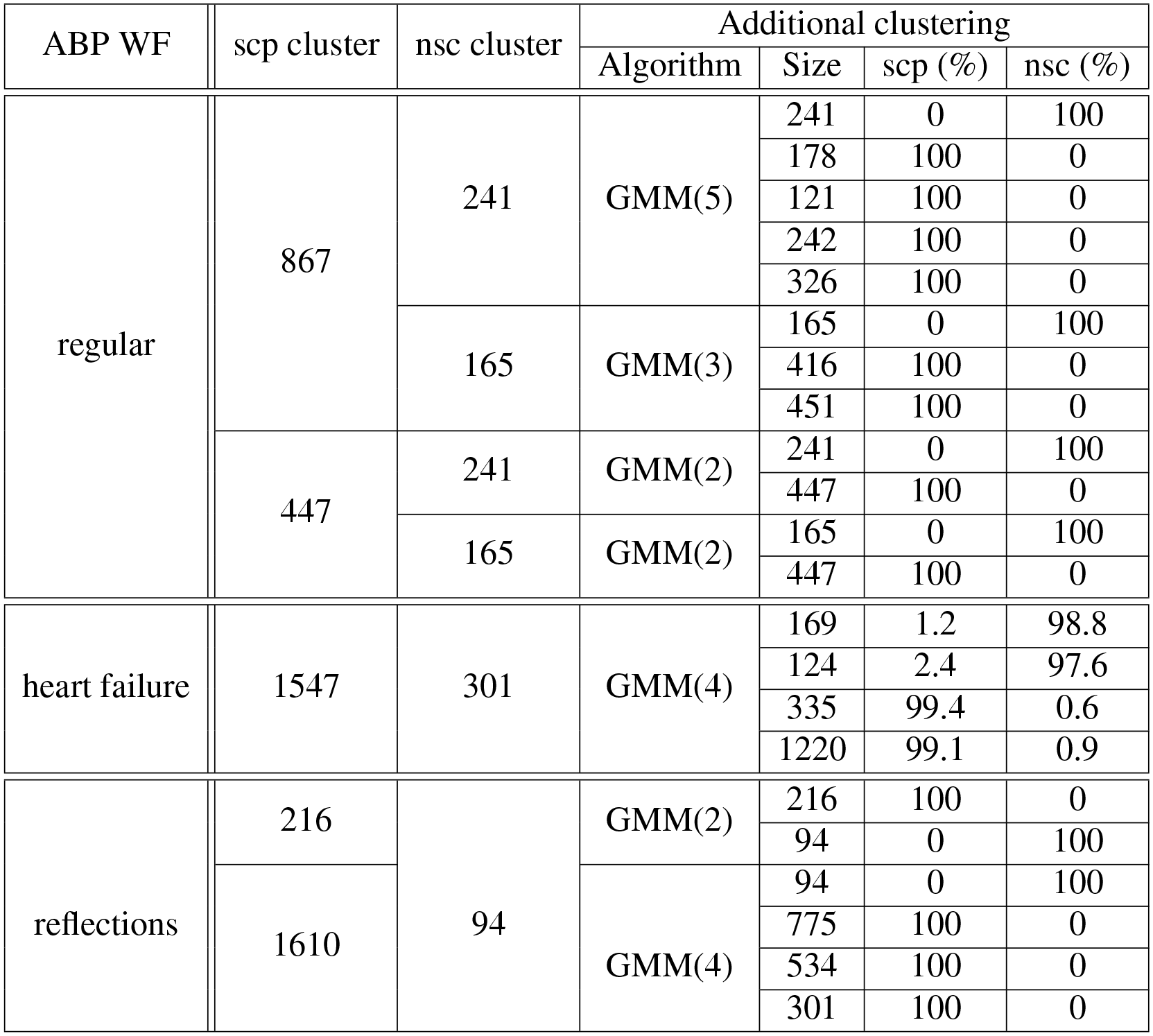
Additional clustering results of pair-wise (scp-nsc) selected ICP clusters corresponding to chosen ABP waveforms.

#### 3.2.3 Classification

The data arranged in Table 4 was exploited as a dataset for end-to-end training and testing the FCN classifier described in Section 2.5.1. The six types of ICP pulses (Fig. 5) that can be assigned to the ABP waveforms regular, heart failure and reflective input, respectively, and separated by sc-index come naturally as class labels. Based on this premise a training dataset was curated (see Table 5). Even though the dataset is highly imbalanced, after fitting the FCN on the training set (80 % of the dataset) it reached an average accuracy of 96 % on the hold-out test set.

**Table 5:** Training dataset consisting of six classes, divided into **scp** and **nsc**, additionally split into regular, heart failure and multiple reflections input ABP waveforms.

| Selected correlation type | input ABP-WF | Nr. of ICP pulses |
| --- | --- | --- |
| scp | regular | 2181 |
|  | heart failure | 1555 |
|  | reflections | 1826 |
| nsc | regular | 406 |
|  | heart failure | 293 |
|  | reflections | 95 |

With the purpose of understanding what determined the model to classify each type of pulses, a part of the test set was subjected to the CAMs technique (see Figs. S6 and S7), as described in Section 2.5.1. One common characteristic among the majority of these CAMs is the weight with which the initial slope of the pulse contributes to its classification. This phenomenon is especially striking in the *scpcommon* class (see first panel in Fig. S6), where the majority of the contribution is focused right at the beginning of the pulse sample, since one would expect that the label will be assigned based on the ripples recorded in 150 *−* 500 segment.

One definite conclusion that can be drawn from these results is that the ABP waveforms, viewed as an input driving the ICP, definitely introduce detectable changes in the shape of ICP waves. Furthermore, these alterations are not substantial enough to smear out differences between sc types. Moreover, the techniques applied, namely PCA for dimensionality reduction and GMMs with HDBSCAN for clustering, proved to be very accurate and practical in uncovering patterns and correlations in the ICP and ABP pulsatile data never observed before.

## 4 Conclusions

The variations induced by a diminished cerebral compliance on ICP waveforms depend on a large number of internal and external influences. For an automatic detection of medically relevant information contained in these variations it is therefore inevitable to categorize the most apparent influences and to reduce disrupting factors. In this study we tried to minimized the subjective influence of a human expert by selecting ICP data segments hallmarked by diminished or intact cerebral compliance via a well established mathematical method called selected correlation analysis (SCA). From this labeled data we extracted the appropriate ICP waveforms for further analysis. Using dimensionality reduction in combination with clustering methods we found multiple modes of pulsatile ICPs originating from the two different states of cerebral compliance under investigation. This multitude of modes could be further exploited by constructing a training data set for a custom fully convolutional network. A small number of training steps were enough for the network to reliably distinguish the two states of pulsatile ICPs.

Subsequently we focused on the influence of ABP waveform variants triggered by various diseases on the ICP waveform sets reflecting the state of the cerebral compliance. Once again using methods identical to those used for ICP, we were able to show that distinct variants of ABP waveforms generate detectable sub-clusters in the ICP modes. But more importantly, these sub-clusters were still separable in terms of cerebral compliance state. Based on these observations we constructed a second dataset that can be used for training a classifier for distinguishing between six different kinds of ICPs - two states of the cerebral compliance times three sub-classes of ABP waveform variants. Our network was trained on this dataset and achieved high accuracy, hence can serve as a baseline for pulsatile ICP classification. These findings provide a proof of concept for further development of this type of analysis targeting the real time detection of numerous cerebral pathophysiologies occurring during traumatic brain injury.

## 5 Limitations

Even though we succeeded in detecting and identifying several hitherto unknown types of ICP and ABP pulses, we still lack decent knowledge on their interpretation and significance in a patient-oriented treatment. Our entire methodology can be drastically improved with further research. For example, we used the exact same pre-processing routine for pulsatile ABP as for ICP, but the former could have very different dynamics which could require a different extraction method. Another drawback is the small number of patients present in our cohort. This is a critical variable that has to be increased, in order to sample the majority of conditions for an unbiased and significant diagnosis. All these points have to be seriously considered in order to arrive at an end-to-end system that can be included in a monitor or alert system at the ICU. Our hope is that medical institutions will begin to collaborate more frequently and, more importantly, share their multivariate recordings with institutions and teams specialized in medical data analysis. This type of collaborations is already bringing higher precision and more confidence in the diagnosis of specific pathologies. And with multi-disciplinary efforts medical system’s effectiveness will only be improved.

## 6 Acknowledgments

We acknowledge the deposition of the manuscript on the bioRxiv preprint server under https://doi.org/10.1101/2020.11.17.381517.

## 7 Supplementary Materials

### 7.1 Supplementary tables

**Table 1:** Physiological variables of involved patients

| ID | Gender | Age | Diagnosis | GCS | GOS (last follow up) |
| --- | --- | --- | --- | --- | --- |
| 1 | m | 60.2 | TBI | 10 | 3 |
| 2 | m | 51.6 | SAH | 8 | 1 |
| 3 | f | 51.6 | SAH | 9 | 3 |
| 4 | m | 65.4 | TBI | 6 | 3 |
| 5 | f | 50.4 | SAH | 14 | 5 |
| 6 | f | 43.4 | SAH | 11 | 1 |
| 7 | m | 36.3 | SAH | 12 | 3 |

*SAH* Subarachnoid hemorrhage; *TBI* traumatic brain injury; *GCS* Glasgow coma scale; *GOS* Glasgow outcome scale.

### 7.2 Supplementary figures

**Figure S1:**
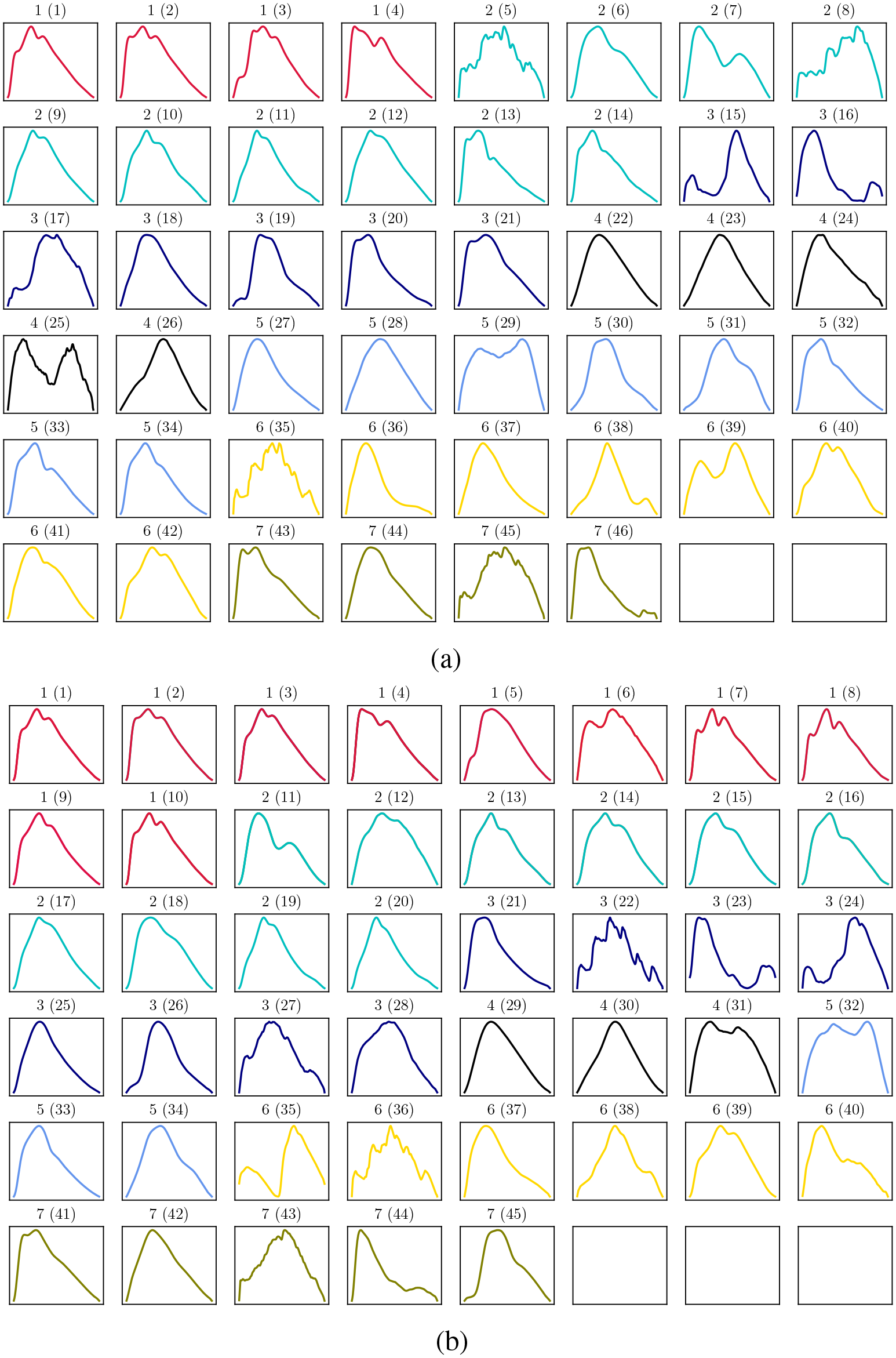
Means of ICP **nsc** (**a**) and **scp** (**b**) clusters. Notation: patient (mean index). Different colours correspond to different patients.

**Figure S2:**
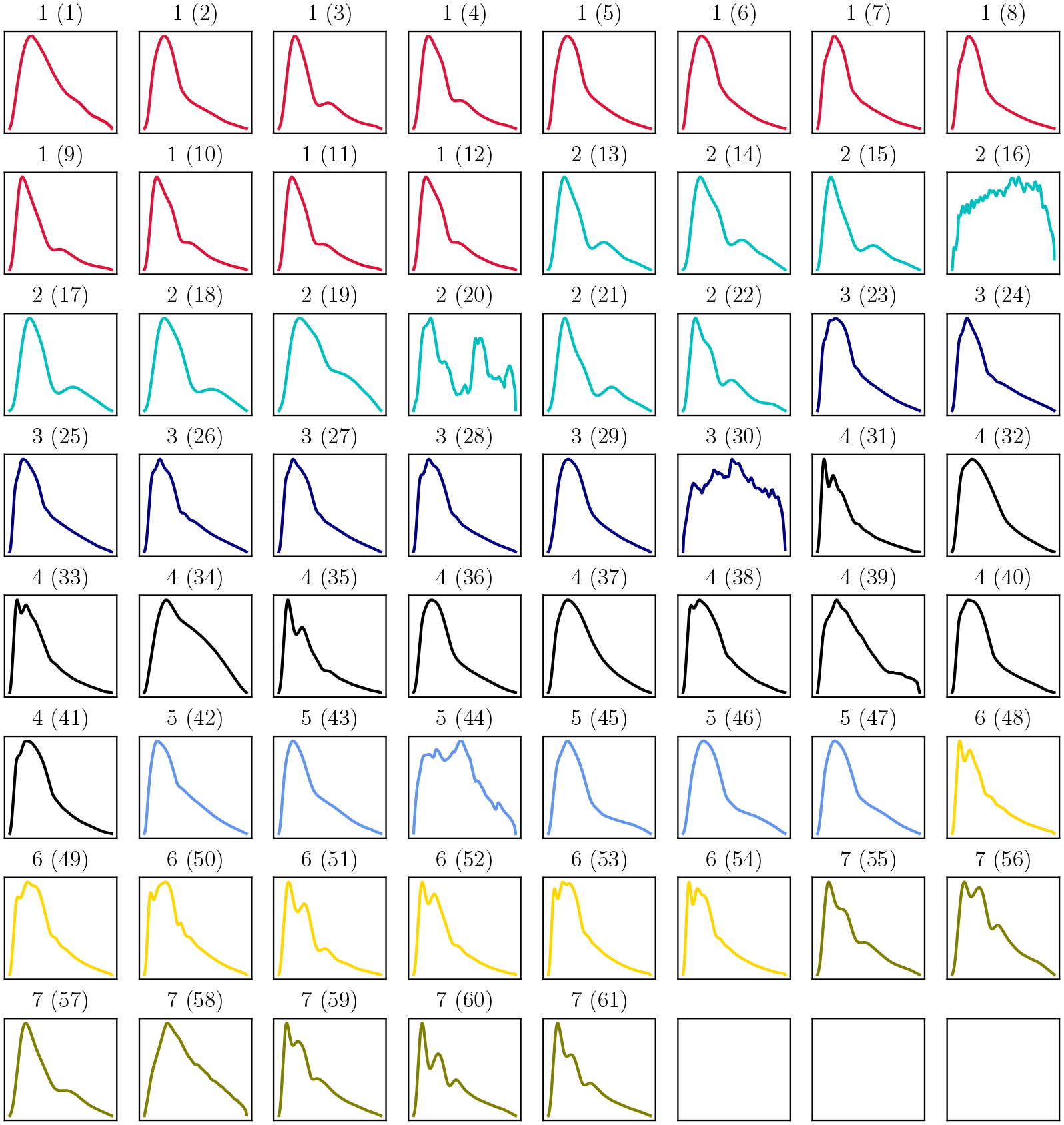
Means of ABP **nsc** clusters. Notation: patient (mean index). Different colours correspond to different patients.

**Figure S3:**
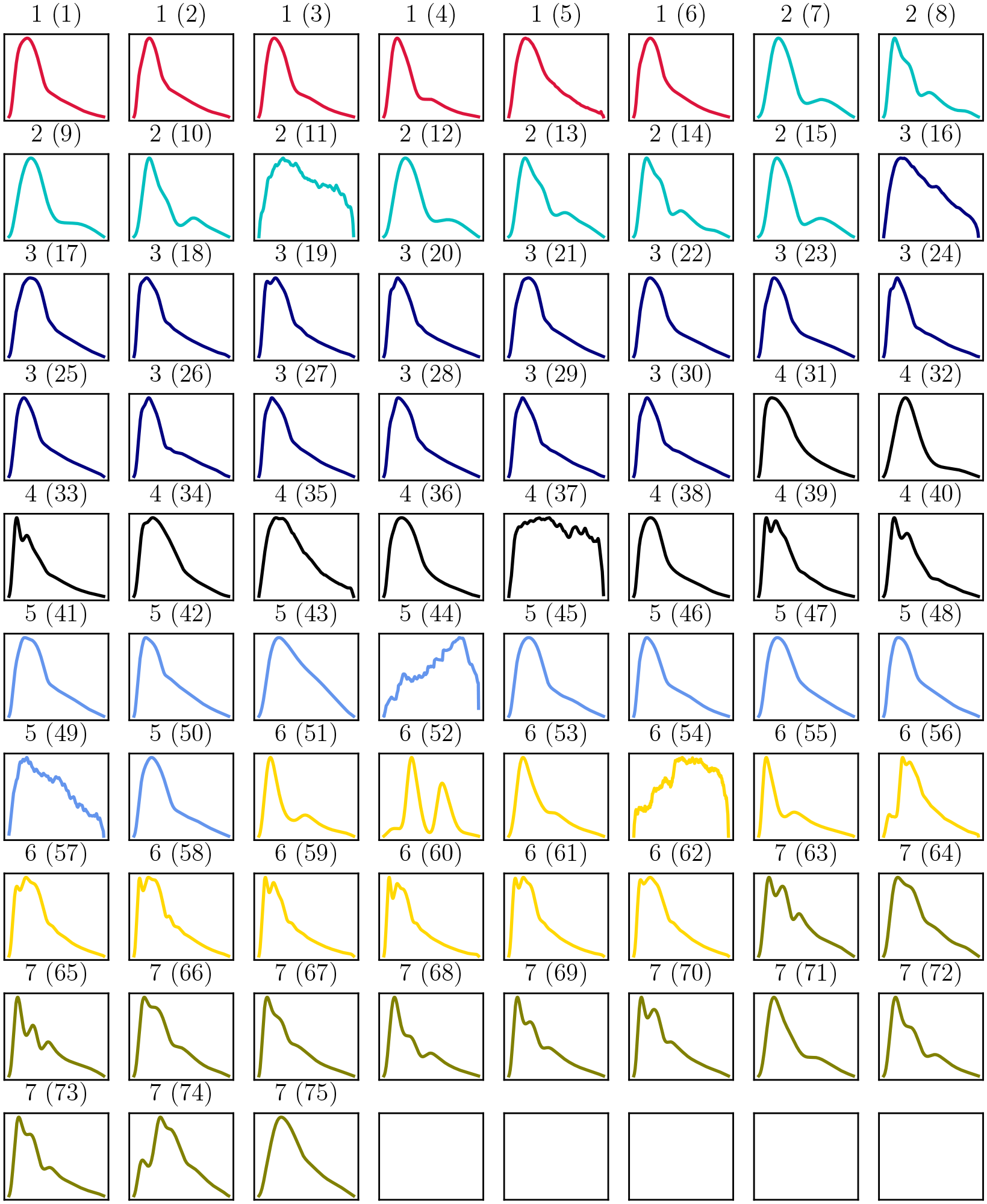
Means of ABP **scp** clusters. Notation: patient (mean index). Different colours correspond to different patients.

**Figure S4:**
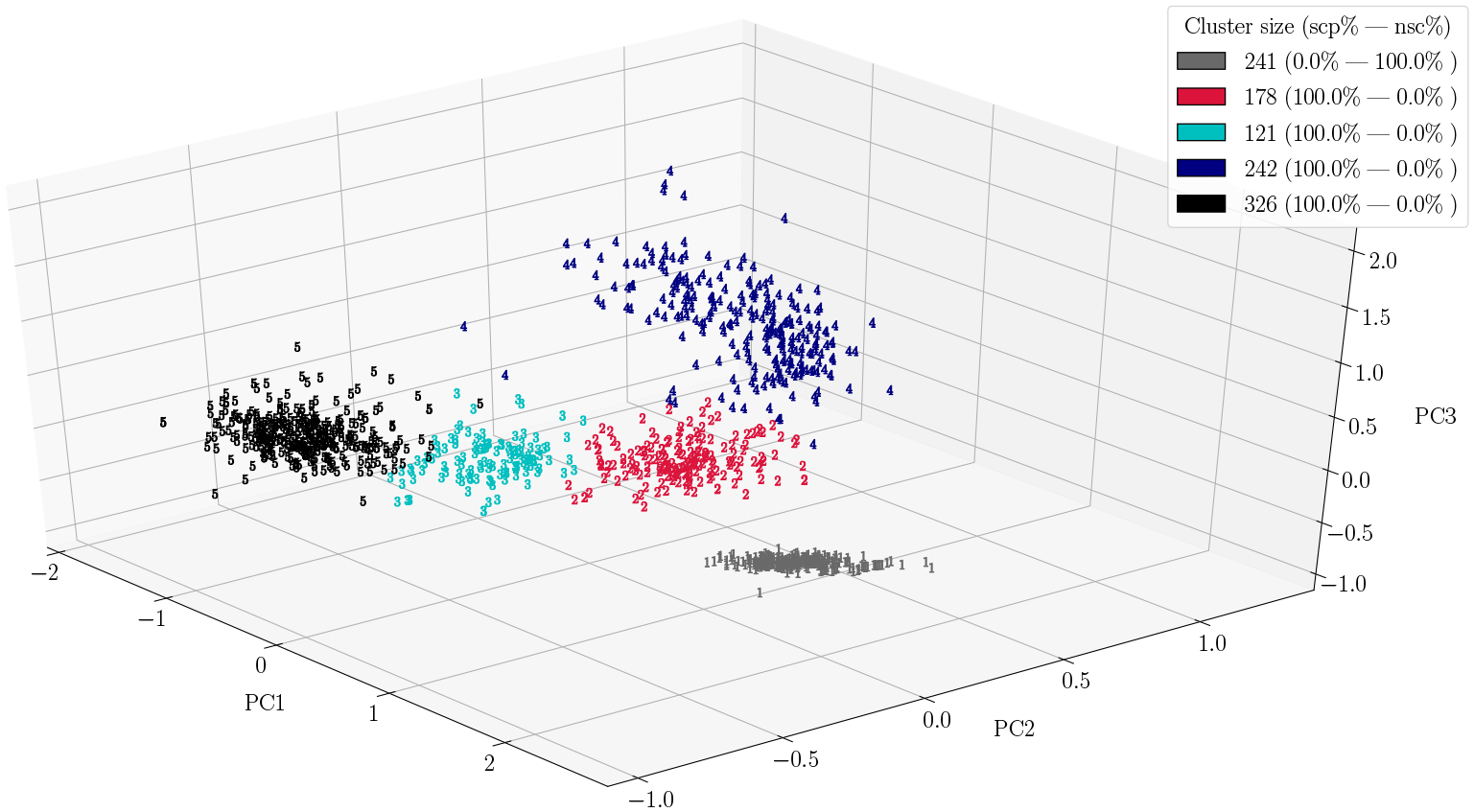
Example of additional clustering using GMMs with 5 components onto the two clusters: 867 **scp** and 241 **nsc** ICP pulses from ABP-WF1.

**Figure S5:**
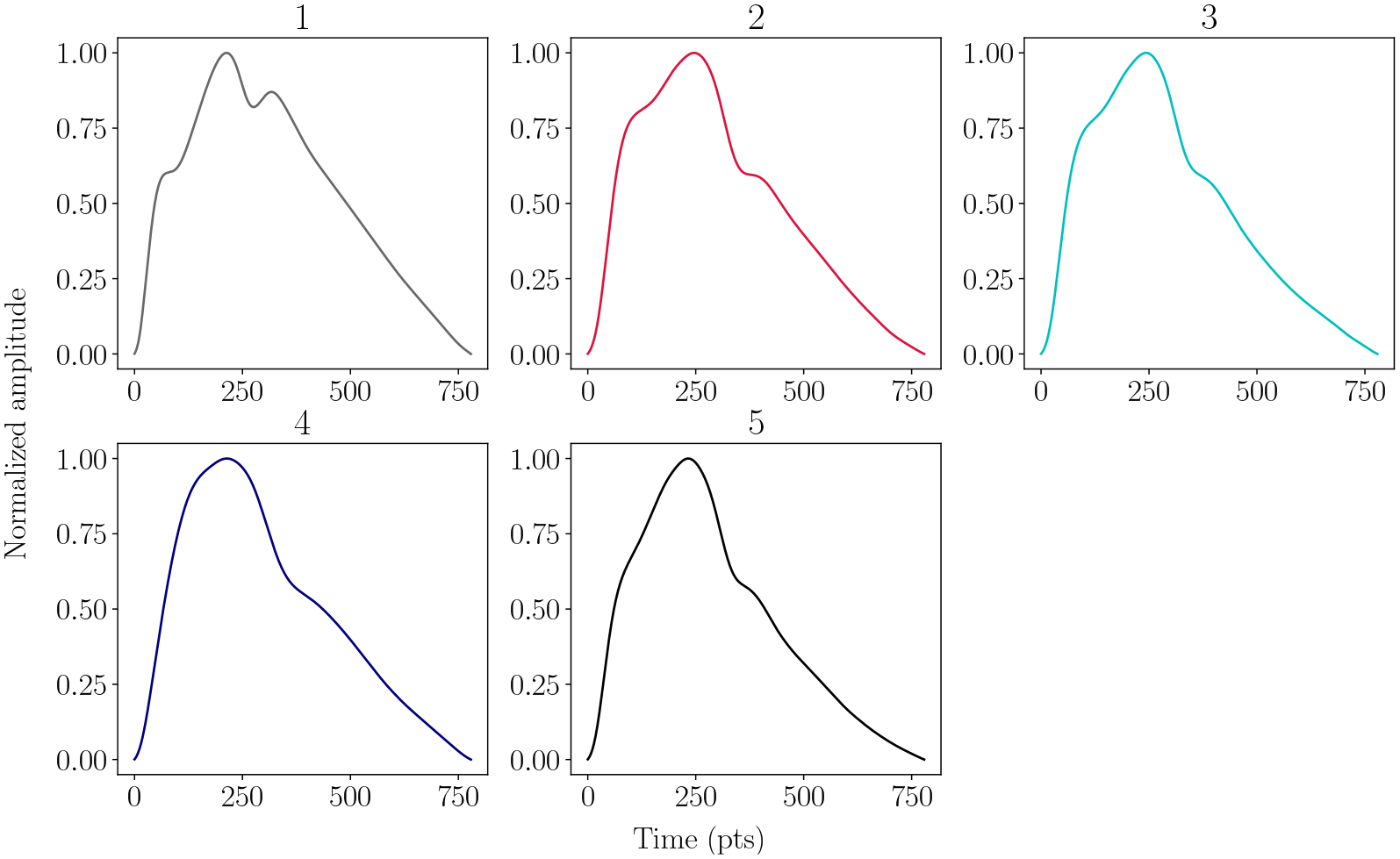
ICP means corresponding to found cluster in the projection (Fig. S4).

### 7.3 Class Activation Maps (CAMs) definition

Given a time series, let *S_k_* (*x*) be the *k*-th activation filter in the last convolutional layer at temporal location *x*. The output of the GAP for filter *k ∈* [1*, M*] is *f_k_* = Σ*_k_ S_k_* (*x*). With 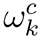 as the weight of the final softmax function for the *k*-th filter and the class *c*, the input of the softmax function will be

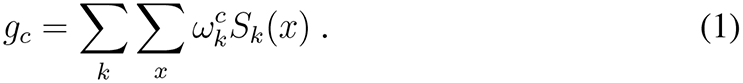

One can define *M_c_* as the class activation map for class *c*, then each temporal element will be given by

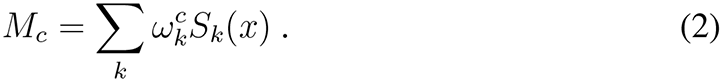

This quantity comprises a univariate time series with each element equal to the weighted sum of the *M* data points at position *x*, the weights being learned by the network [1]. Therefore *M_c_*(*x, y*) will indicate the importance of the activation at location *x_i_* that lead to the classification of the series *y* to class *c*.

**Figure S6:**
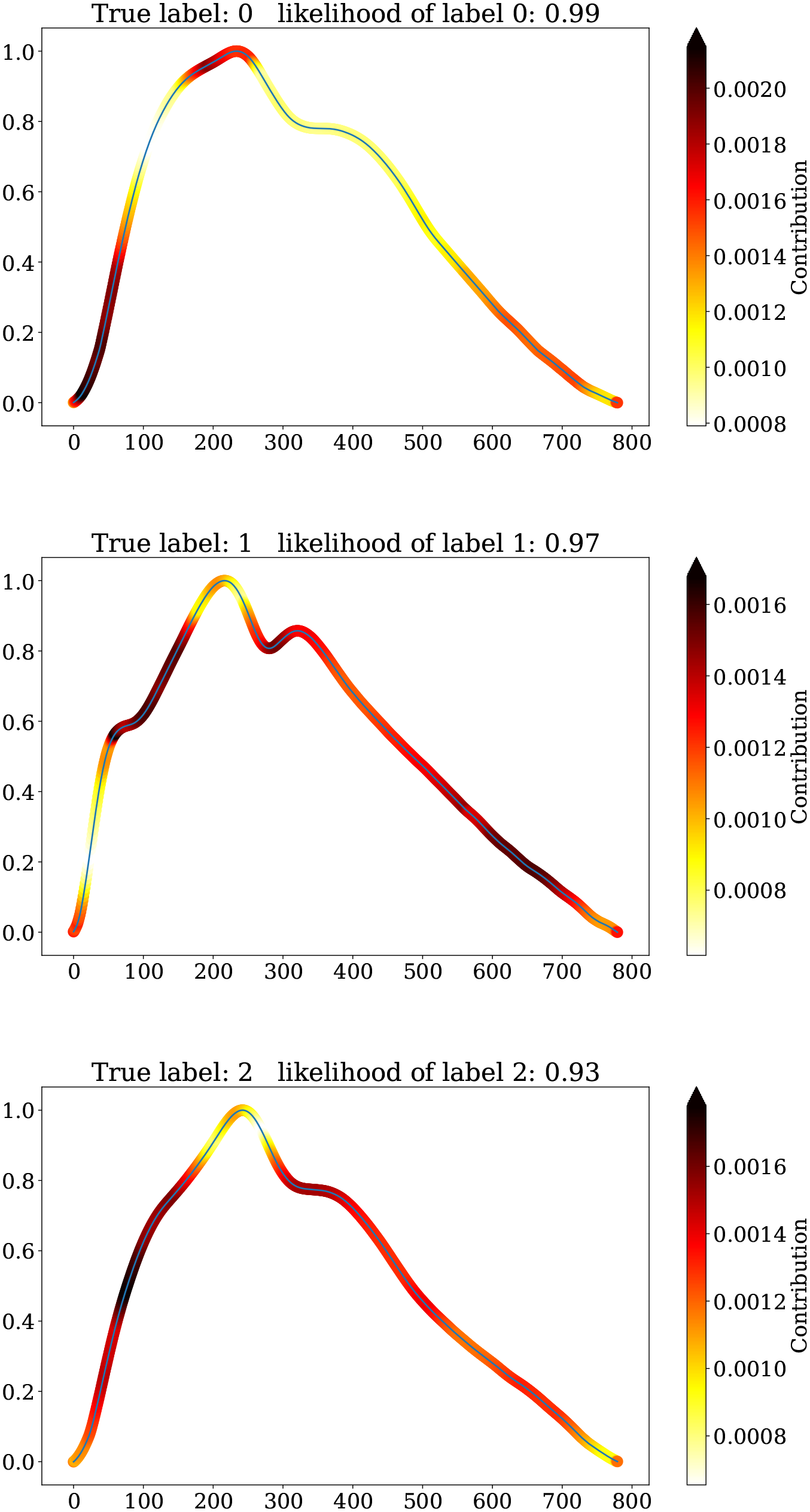
Examples of CAMs from classes 0 2. The true label is displayed together with the predicted one and its `softmax` probability. Darker colors along the shape of the pulses show the regions that contributed the most to the classification.

**Figure S7:**
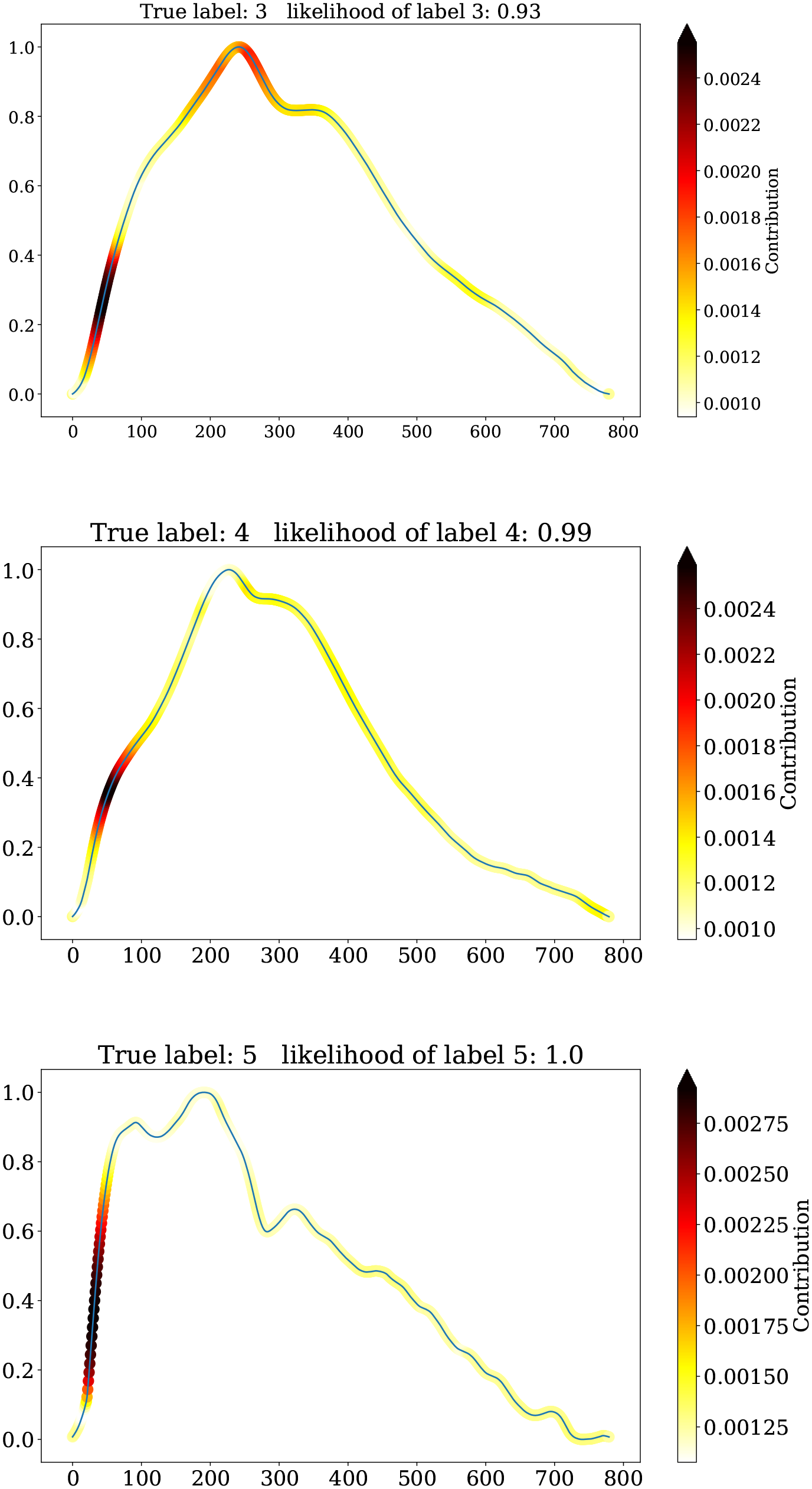
Examples of CAMs from classes 3 → 5. The true label is displayed together with the predicted one and its `softmax` probability. Darker colors along the shape of the pulses show the regions that contributed the most to the classification.

## References

1. Achhammer, T. *An Empirical Mode Decomposition to investigate Intracranial Pressure Profiles to distinguish brain disease states* (Unpublished). MS thesis (Universität Regensburg, Regensburg, Germany, 2017).

2. Bishop, C. M. *Pattern Recognition and Machine Learning* (ed Jordan M. Kleinberg J., S. B.) (Springer-Verlag New York, 2006).

3. Böhm, M., Faltermeier, R., Brawanski, A. & Lang, E. W. Mathematical modeling of human brain physiological data. Physical Review E - Statistical, Nonlinear, and Soft Matter Physics 88 (2013).

4. Campello, R. J., Moulavi, D. & Sander, J. Density-based clustering based on hierarchical density estimates. Lecture Notes in Computer Science (including subseries Lecture Notes in Artificial Intelligence and Lecture Notes in Bioinformatics) 7819 LNAI, 160–172 (2013).

5. Chollet, F. et al. Keras https://keras.io. 2015.

6. Denardo, S. J., Nandyala, R., Freeman, G. L., Pierce, G. L. & Nichols, W. W. Pulse wave analysis of the aortic pressure waveform in severe left ventricular systolic dysfunction. Circulation: Heart Failure 3, 149–156 (2010).

7. Faltermeier, R., Proescholdt, M. A., Bele, S. & Brawanski, A. Parameter Optimization for Selected Correlation Analysis of Intracranial Pathophysiology. Computational and Mathematical Methods in Medicine 2015 (2015).

8. Faltermeier, R., Proescholdt, M. A., Bele, S. & Brawanski, A. Windowed multitaper correlation analysis of multimodal brain monitoring parameters. Computational and Mathematical Methods in Medicine 2015 (2015).

9. Hu, X., Glenn, T., Scalzo, F., Bergsneider, M., Sarkiss, C., Martin, N. & Vespa, P. Intracranial pressure pulse morphological features improved detection of decreased cerebral blood flow. Physiological Measurement (2010).

10. Hu, X., Xu, P., Scalzo, F., Vespa, P. & Bergsneider, M. Morphological clustering and analysis of continuous intracranial pressure. IEEE Transactions on Biomedical Engineering (2009).

11. Ismail Fawaz, H., Forestier, G., Weber, J., Idoumghar, L. & Muller, P. A. Deep learning for time series classification: a review. Data Mining and Knowledge Discovery 33, 917–963. arXiv: 1809.04356 (2019).

12. Jolliffe, I. T. Principal Component Analysis, Second Edition. Encyclopedia of Statistics in Behavioral Science (2002).

13. Jung, A. Statistical analysis of biomedical data. eprint: 10.5283/epub. 10168 (2003).

14. Lin, M., Chen, Q. & Yan, S. Network In Network. arXiv: 1312.4400 (Dec. 2013).

15. Long, J., Shelhamer, E. & Darrell, T. Fully Convolutional Networks for Semantic Segmentation. arXiv: 1411.4038 (Nov. 2014).

16. McInnes, L. & Healy, J. Accelerated Hierarchical Density Based Clustering. *IEEE International Conference on Data Mining Workshops*, ICDMW 2017-Novem, 33–42. arXiv: 1705.07321 (2017).

17. Nirmalan, M. & Dark, P. M. Broader applications of arterial pressure wave form analysis. *Continuing Education in Anaesthesia*, Critical Care and Pain 14, 285–290 (2014).

18. Nucci, C. G., De Bonis, P., Mangiola, A., Santini, P., Sciandrone, M., Risi, A. & Anile, C. Intracranial pressure wave morphological classification: automated analysis and clinical validation. Acta Neurochirurgica 158, 581– 588 (2016).

19. Proescholdt, M. A., Faltermeier, R., Bele, S. & Brawanski, A. Detection of impaired cerebral autoregulation using selected correlation analysis: A validation study. Computational and Mathematical Methods in Medicine 2017 (2017).

20. Wang, Z., Yan, W. & Oates, T. Time Series Classification from Scratch with Deep Neural Networks: A Strong Baseline. CoRR abs/1611.0. arXiv: 1611.06455 (2016).

21. Zeiler, A. Weighted Sliding Empirical Mode Decomposition and its Application to Neuromonitoring Data. eprint: 10.5283/epub.27783 (2012).

22. Zhou, B., Khosla, A., Lapedriza, A., Oliva, A. & Torralba, A. Learning Deep Features for Discriminative Localization. Proceedings of the IEEE Computer Society Conference on Computer Vision and Pattern Recognition, 2921–2929. arXiv: 1512.04150 (2016).

## References

1. Ismail Fawaz, H., Forestier, G., Weber, J., Idoumghar, L. & Muller, P. A. Deep learning for time series classification: a review. Data Mining and Knowledge Discovery 33, 917–963. arXiv: 1809.04356 (2019).

